# Chemoproteomic profiling of substrate specificity in gut microbiota-associated bile salt hydrolases

**DOI:** 10.1101/2024.04.01.587558

**Authors:** Lin Han, Augustus Pendleton, Adarsh Singh, Raymond Xu, Samantha A. Scott, Jaymee A. Palma, Peter Diebold, Kien P. Malarney, Ilana L. Brito, Pamela V. Chang

## Abstract

The gut microbiome possesses numerous biochemical enzymes that biosynthesize metabolites that impact human health. Bile acids comprise a diverse collection of metabolites that have important roles in metabolism and immunity. The gut microbiota-associated enzyme that is responsible for the gateway reaction in bile acid metabolism is bile salt hydrolase (BSH), which controls the host’s overall bile acid pool. Despite the critical role of these enzymes, the ability to profile their activities and substrate preferences remains challenging due to the complexity of the gut microbiota, whose metaproteome includes an immense diversity of protein classes. Using a systems biochemistry approach employing activity-based probes, we have identified gut microbiota-associated BSHs that exhibit distinct substrate preferences, revealing that different microbes contribute to the diversity of the host bile acid pool. We envision that this chemoproteomic approach will reveal how secondary bile acid metabolism controlled by BSHs contributes to the etiology of various inflammatory diseases.

## Introduction

Systems biochemistry is an emerging framework that combines the scalability of systems biology approaches and the precision of mechanistic enzymology to ascribe function to previously uncharacterized enzymes on the proteome-wide scale (Ali and Bar-Peled, 2024; Sung et al., 2020). Such approaches have been immensely useful for tapping into the dark matter of mammalian proteomes, as the activities of a significant number of proteins remain unknown. The metagenome of the human gut microbiome, which comprises hundreds of trillions of microorganisms including bacteria, viruses, fungi, and parasites, is predicted to encode at least 170 million proteins, of which approximately 40% lack functional annotations (Almeida et al., 2021). This important microbial ecosystem greatly impacts the host through regulation of metabolism, immunity, and inflammation in many (patho)physiological contexts (Ansaldo et al., 2021; Fan and Pedersen, 2021; Kayama et al., 2020).

Despite the significance of the gut microbiome in human health and disease, mechanistic understanding of the myriad functions of the microbiota remains a grand challenge, including their metabolic and biosynthetic potential (Koppel and Balskus, 2016). The gut microbiota harbor a vast multitude of proteins involved in biochemical transformations of important secondary metabolites (Dorrestein et al., 2014; McCarville et al., 2020; Skelly et al., 2019); yet, the activities of these enzyme families within the complex microbial milieu remain poorly understood due to a lack of tools to characterize their functions on a global scale. Because of the sheer number of putative enzymatic activities encoded by the gut microbiome, the use of traditional reductionist approaches to comprehensively characterize the diverse biochemistry exhibited by entire enzyme classes would be highly inefficient. Thus, we posit that emerging systems biochemistry strategies will be needed to elucidate protein activities and functions within the gut metaproteome.

Bile acids are amphipathic metabolites whose primary role is to aid in the solubilization of dietary lipophilic nutrients in the intestines, where they are further modified by gut microbiota-associated enzymes to form a wide variety of secondary metabolites (Ridlon et al., 2006). Host- and microbially-produced bile acids act as signaling molecules within the host, where they activate several nuclear and G protein-coupled receptors, including farnesoid X receptor (FXR), vitamin D receptor (VDR), pregnane X receptor (PXR), constitutive androstane receptor (CAR), and G protein-coupled bile acid receptor 1 (GPBAR1 or TGR5) to modulate lipid and glucose homeostasis, xenobiotic metabolism, and immunity (Cai et al., 2022; Collins et al., 2023; Perino and Schoonjans, 2022). Due to their effects on these physiological processes, bile acids impact many inflammatory diseases, including infection, cancer, inflammatory bowel diseases, and metabolic syndrome. These metabolites also shape microbial composition within the gut because their antimicrobial properties are toxic to certain bacteria (Begley et al., 2005).

Bile acids are initially synthesized by host enzymes within the liver from cholesterol into two primary bile acids, cholic acid and chenodeoxycholic acid (Ridlon et al., 2006). These bile acids are then conjugated with glycine and taurine by host enzymes in the liver to form glyco- and tauro-conjugated bile acids. These metabolites are further modified by the gut microbiota when they are postprandially secreted into the intestines to aid in digestion of lipophilic nutrients. Such transformations include deconjugation, oxidation, epimerization, and dehydroxylation by microbial enzymes that ultimately convert primary bile acids into secondary bile acids, including deoxycholic acid and lithocholic acid. Recent reports have demonstrated that primary bile acids can also be reconjugated by gut bacteria with many proteinogenic amino acids, including leucine, phenylalanine, and tyrosine, which greatly expands the pool of bile acids within the host (Garcia et al., 2022; Gentry et al., 2024; Guzior et al., 2024; Lucas et al., 2021; Mohanty et al., 2024; Neugebauer et al., 2022; Quinn et al., 2020; Rimal et al., 2024; Shalon et al., 2023).

Bile salt hydrolases (BSH, EC 3.5.1.24) are cysteine hydrolases expressed exclusively by the gut microbiota that are responsible for the deconjugation of conjugated BAs to release the amino acid and the free bile acid (Figure 1A) (Foley et al., 2019). BSHs have been termed gatekeeper enzymes in secondary bile acid metabolism because they catalyze the gateway reaction that is thought to be required for all subsequent biochemical transformations by the host or gut microbiota (Begley et al., 2006). Interestingly, recent work suggests that BSHs are responsible for reconjugation of amino acids to deconjugated bile acids because they also act as N-acyl aminotransferases, which indicates that our understanding of intestinal bile acid metabolism is still evolving (Guzior et al., 2024; Rimal et al., 2024). Accordingly, BSHs remain a critical enzyme class in secondary bile acid metabolism because they control the overall composition of bile acid pool, which ultimately influences gut microbial composition and activation of specific host signaling pathways. Despite the importance of this hydrolase family in many host physiological processes, few tools exist to probe their activity and substrate preference due to the multitude of distinct BSH enzymes within the gut microbiota and the number of possible bile acid substrates for each bacterial BSH.

**Figure 1.**
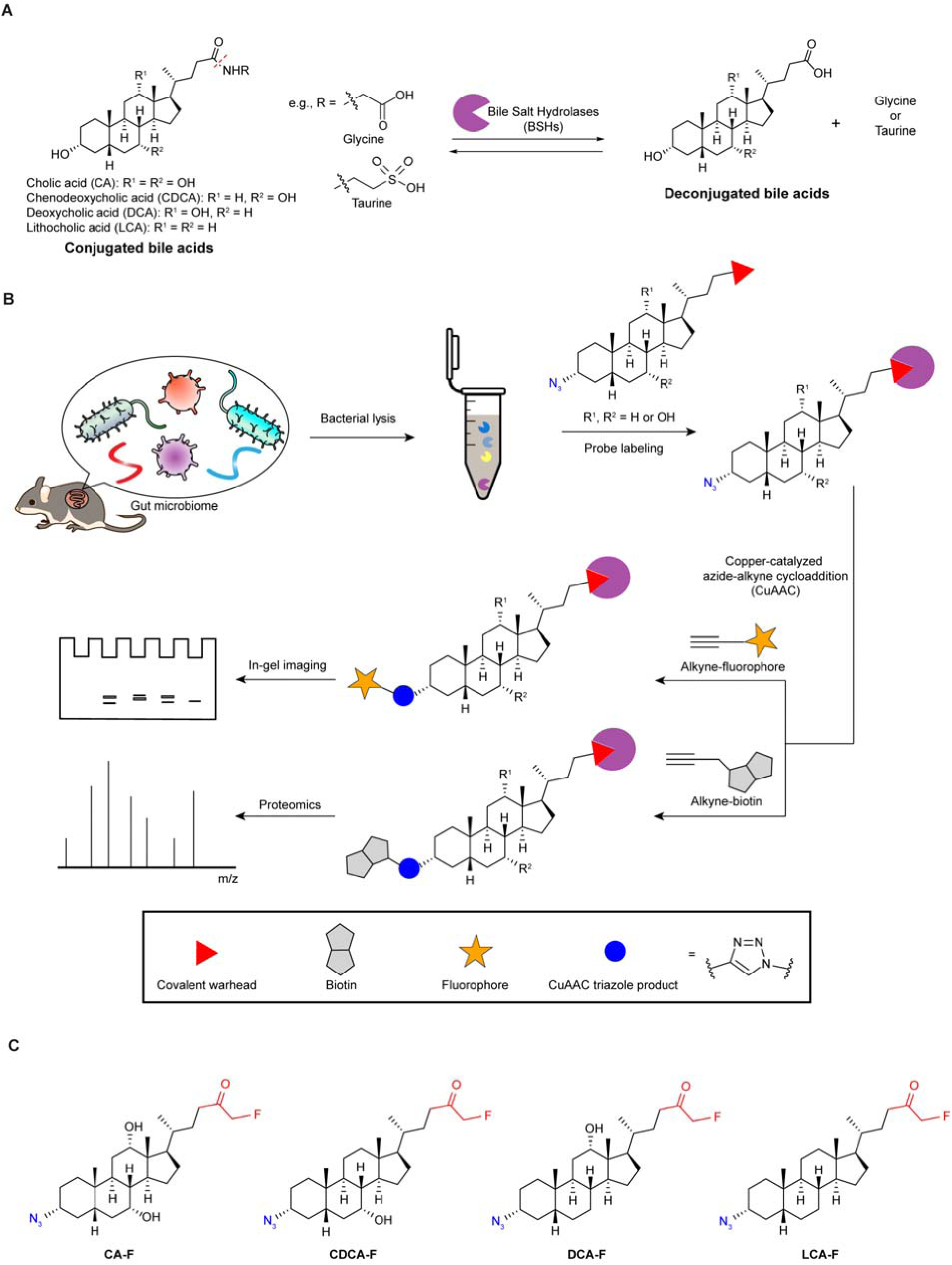
Chemoproteomic, activity-based approach for profiling bile salt hydrolase (BSH) activity within the gut microbiome. (A) BSH carries out the deconjugation reaction of glyco- and tauro-conjugated bile acids, which is the first major step of secondary bile acid metabolism in the intestines, and is also capable of carrying out the reconjugation of a wide variety of amino acids. (B) Scheme of overall chemical strategy, termed **BSH**-**T**agging and **R**etrieval with **A**ctivity-based **P**robes (BSH-TRAP), to covalently label active BSH enzymes via the active site cysteine residue using activity-based probes and Cu-catalyzed azide-alkyne cycloaddition (CuAAC) click chemistry to tag labeled enzymes with an affinity handle or contrast agent for pulldown or imaging of BSH activity. (C) Structures of the panel of fluoromethylketone (FMK) activity-based probes used to identify BSH activity in the gut microbiome in this study.

Thus, as traditional biochemical approaches to characterize enzymatic activity are limited in their ability to probe biochemical activity on the systems level, we have developed a chemoproteomic platform, **B**ile **S**alt **H**ydrolase-**T**agging and **R**etrieval with **A**ctivity-based **P**robes (BSH-TRAP, Figure 1B), to profile the activities of BSHs within complex biological settings, such as the gut microbiome (Parasar and Chang, 2022; Parasar et al., 2019). This activity-based protein profiling (ABPP) approach utilizes tailored activity-based probes (ABPs) that covalently label the catalytic cysteine residue within the BSH enzyme active site. ABPs typically comprise a targeting group to direct the molecule to the proteins of interest, a covalent warhead or electrophilic group for trapping reactive amino acid residues near the probe binding site, and a click chemistry handle (e.g., azide or alkyne) that enables subsequent labeling for visualization or identification (Niphakis and Cravatt, 2014; Sanman and Bogyo, 2014).

Previously, we developed a first-generation BSH ABP based on cholic acid, a common BSH substrate, that contains an acyloxymethylketone (AOMK) warhead, which is selective for cysteine residues, and an azide that can undergo secondary labeling with an alkyne containing probe using Cu(I)-catalyzed azide-alkyne cycloaddition (CuAAC) (Parasar et al., 2019). Using this cholic acid-based ABP, we profiled BSH activities within gut microbes and identified active BSH enzymes within healthy and colitic mouse gut microbiomes. Additional approaches have been developed using chemical probes containing a fluorogenic or luminogenic group to monitor BSH activity (Brandvold et al., 2019; Khodakivskyi et al., 2021; Kombala et al., 2023; Sveistyte et al., 2020), alternative covalent warheads to profile and inhibit BSH activity (Adhikari et al., 2020; Brandvold et al., 2021), and photoaffinity labels to identify bile acid-binding proteins (Forster et al., 2022; Liu et al., 2022; Yang et al., 2022, 2023; Zhuang et al., 2017). Despite the number of emerging chemical tools to study BSH activities, the substrate preferences within the BSH family at a metaproteome level remain unknown.

Here, we sought to address this question by profiling the substrate specificity of gut microbiota-associated BSHs using a broadened version of the BSH-TRAP strategy that leverages a new panel of chemical probes that comprise analogs of different BSH substrates based on the bile acid steroid core. This toolkit contains ABPs for both prominent human primary bile acids, cholic acid (CA) and chenodeoxycholic acid (CDCA), and secondary bile acids, deoxycholic acid (DCA) and lithocholic acid (LCA), (Figure 1C, Scheme S1, Data S1). These probes bear an azido click chemistry handle and a fluoromethylketone (FMK) warhead, which like AOMK also targets cysteine residues but is more sensitive than the AOMK group due to its increased reactivity (Kam et al., 2004; Sanman and Bogyo, 2014). As our previous BSH ABP was based on CA, we hypothesized that this expanded chemical toolkit comprising these four probes, termed CA-F, CDCA-F, DCA-F, and LCA-F (Figure 1C), would allow us to globally characterize BSH substrate preferences for the bile acid steroid core within the gut microbiome using a systems biochemistry approach.

## Results

### Panel of BSH ABPs labels BSH via the catalytic cysteine residue

We first evaluated whether the panel of BSH ABPs (Figure 1C) could label active BSH in vitro. We expressed wild-type *Clostridium perfringens* BSH, also known as choloylglycine hydrolase (CGH, EC 3.5.1.24), in *Escherichia coli*, followed by purification as previously described (Malarney and Chang, 2023; Parasar et al., 2019). To assess whether the probes could covalently label the enzyme, we treated purified CGH with CA-F, CDCA-F, DCA-F, or LCA-F for various amounts of time (Figure 2A, C, E, G, Figure S1A, C, E, G), followed by CuAAC tagging with a Rhodamine 110-alkyne conjugate (Fluor 488-alkyne). Analysis by gel electrophoresis and in-gel fluorescence imaging found that the labeling increased over 10 min and reached a maximum by 30 min. As a negative control, we also generated the catalytically dead Cys2Ser (C2S) CGH mutant and found that the probes did not label this enzyme (Parasar et al., 2019), which demonstrates the specificity of the labeling for the catalytic cysteine (Figure 2A, C, E, G, Figure S1A, C, E, G). We also demonstrated that the probes labeled wild-type CGH in a dose-dependent manner, with detectable labeling at 100 nM that saturated at 1 μM (Figure 2B, D, F, H, Figure S1B, D, F, H). These results indicate that the panel of BSH ABPs covalently label catalytic active site cysteine residues within BSH.

**Figure 2.**
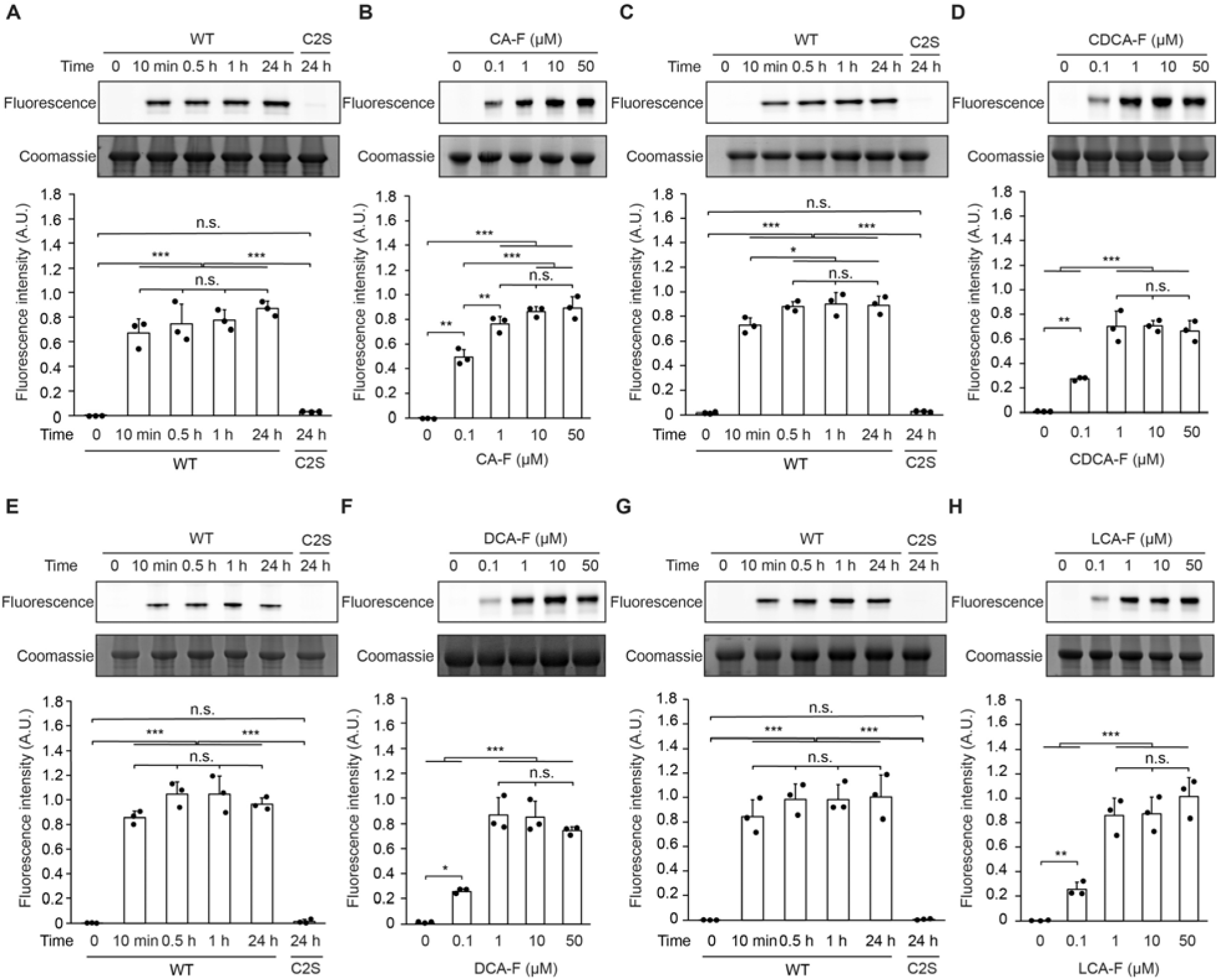
ABPs label *Clostridium perfringens* BSH in a time- and dose-dependent manner. Wild-type (WT) or mutant (Cys2Ser, C2S) BSH from *C. perfringens* was treated with (A) CA-F, (C) CDCA-F, (E) DCA-F, and (G) LCA-F (10 µM) for various amounts of time (indicated) or (B, D, F, H) varying concentrations of probes (indicated) at 37 °C for 1 h, after which the samples were tagged using the copper-catalyzed azide-alkyne cycloaddition (CuAAC) with Fluor 488-alkyne. The samples were analyzed by SDS-PAGE and visualized by in-gel fluorescence. The bands were quantified by densitometry using ImageJ (bottom panels). A.U. = arbitrary unit. Error bars represent standard deviation from the mean. One-way ANOVA followed by post hoc Tukey’s test: * p<0.05, ** p< 0.01, *** p<0.001, n.s. = not significant, n = 3.

### BSH ABPs label endogenous BSH within model gut anaerobic bacteria

We next determined whether the panel of probes could label endogenously expressed BSH from two human gut anaerobes, *Bacteroides fragilis* and *Bifidobacterium longum*. Lysates from these bacteria were labeled with CA-F, CDCA-F, DCA-F, or LCA-F for 1 h, followed by CuAAC tagging with Fluor 488-alkyne and the in-gel fluorescence assay. We found that the panel of probes labeled a protein band at the expected molecular weight (35-37 kDa) for the BSH monomers expressed by these bacteria (Figure 3, Figure S2) (Cerdeño-Tárraga et al., 2005; Sela et al., 2008). When we pre-treated the lysates with iodoacetamide, the probe labeling was eliminated, validating that the labeling is cysteine-selective.

**Figure 3.**
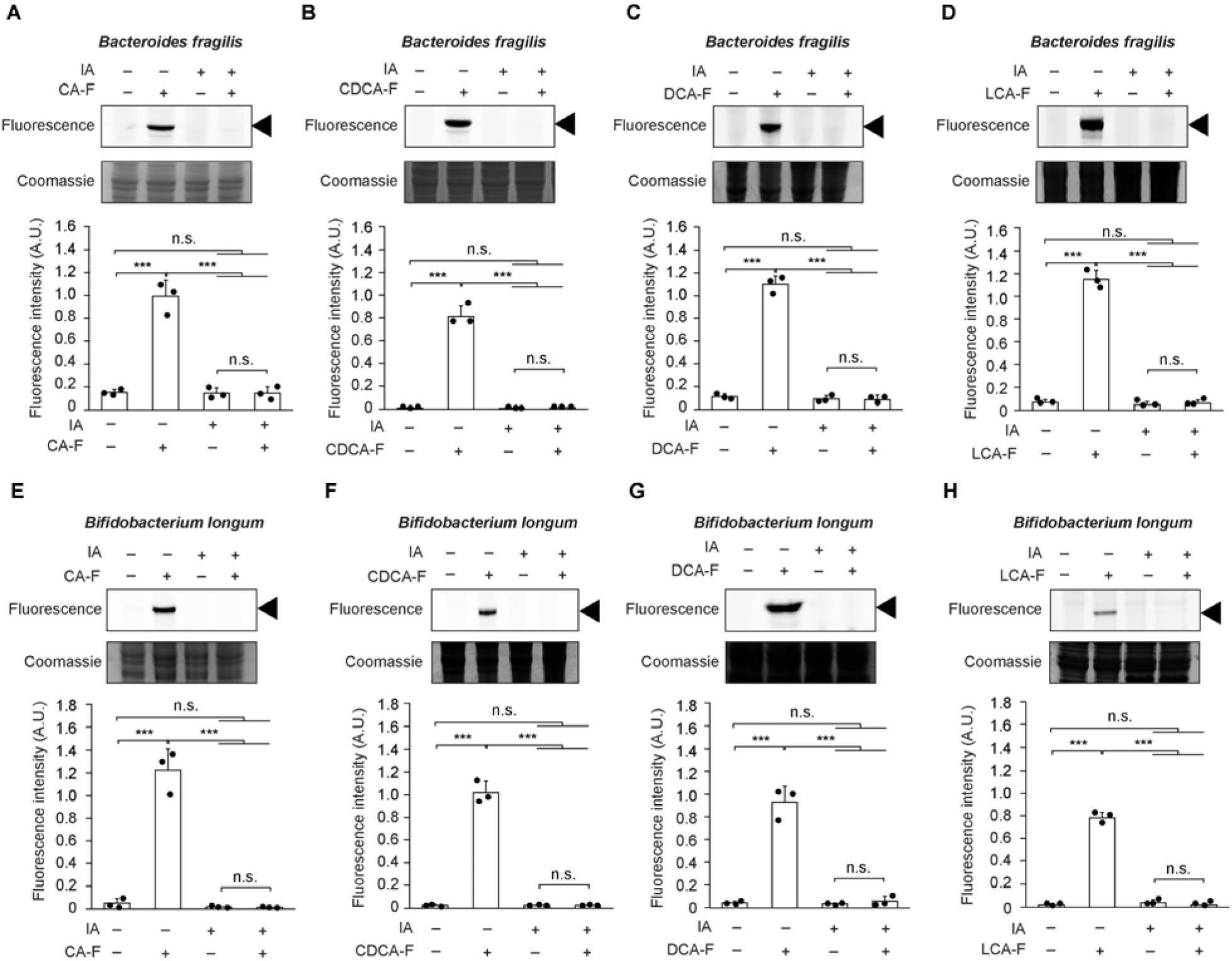
ABPs label BSH from human gut anaerobes. ABPs (indicated, 10 µM) were incubated with lysates from (A-D) *Bacteroides fragilis*, or (E-H) *Bifidobacterium longum* at 37°C for 1 h. Lysates were treated with iodoacetamide (IA, 50 mM) prior to incubation with probes as a negative control. Following probe labeling, CuAAC tagging was carried out with Fluor 488-alkyne. Samples were subjected to SDS-PAGE, and the gel was visualized using fluorescence, followed by Coomassie staining. Arrowhead indicates BSH monomers at 35-37 kDa. The bands were quantified by densitometry using ImageJ (bottom panels). A.U. = arbitrary unit. Error bars represent standard deviation from the mean. One-way ANOVA followed by post hoc Tukey’s test: *** p<0.001, n.s. = not significant, n = 3.

We further demonstrated that the panel of ABPs could be used to determine BSH substrate specificity by labeling *B. fragilis* (Figure 4A, S3A) and *B. longum* (Figure 4B, S3B) with CA-F, CDCA-F, DCA-F, or LCA-F for 10 min, followed by CuAAC tagging with Fluor 488-alkyne and the in-gel fluorescence assay. We found that CA and LCA were the least preferred bile acids for *B. fragilis* and *B. longum*, respectively, whereas the preference for the remaining bile acids were comparable. Importantly, these results are similar to existing in vitro biochemical studies with these individual BSHs (Song et al., 2019). Thus, our panel of probes can, in a single experiment using a highly complex sample (i.e., a cell lysate), reveal the substrate specificity of BSHs endogenously expressed within relevant human gut bacteria.

**Figure 4.**
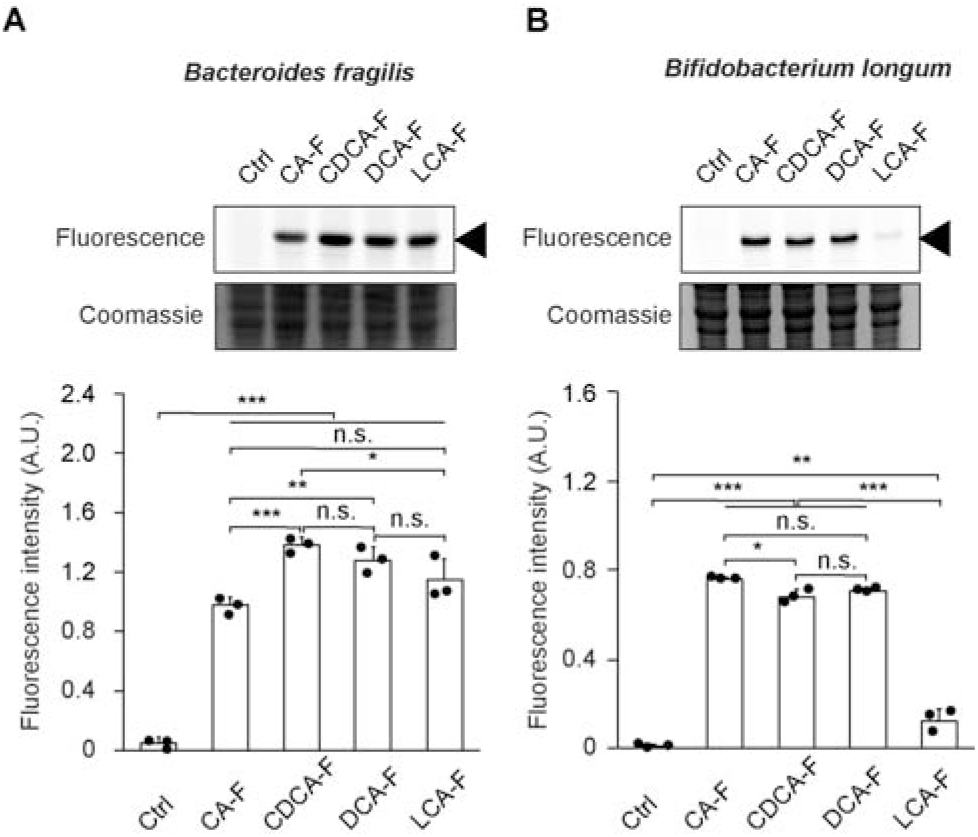
ABPs reveal BSH substrate specificity in human gut anaerobes. ABPs (indicated, 50 µM) or vehicle (DMSO, Ctrl) were incubated with lysates from (A) *Bacteroides fragilis* or (B) *Bifidobacterium longum* at 37 °C for 10 min. Following probe labeling, CuAAC tagging was carried out with Fluor 488-alkyne. Samples were subjected to SDS-PAGE, and the gel was visualized using fluorescence, followed by Coomassie staining. Arrowhead indicates BSH monomers at 35-37 kDa. The bands were quantified by densitometry using ImageJ (bottom panels). A.U. = arbitrary unit. Error bars represent standard deviation from the mean. One-way ANOVA followed by post hoc Tukey’s test: * p<0.05, ** p< 0.01, *** p<0.001, n.s. = not significant, n = 3.

### Panel of BSH ABPs can enrich for endogenous BSH from human gut bacteria

As BSH-TRAP is designed to enable identification of BSHs from human gut anaerobes, we applied the panel of probes to enrich for active BSHs from *B. fragilis* (Figure 5A-D, S4A-D) and *B. longum* (Figure 5E-H, S4E-H). Bacterial lysates were treated with CA-F, CDCA-F, DCA-F, or LCA-F for 1 h, followed by CuAAC tagging with a biotin-alkyne reagent. Labeled proteins were then affinity enriched with streptavidin agarose, and the samples were analyzed by gel electrophoresis, followed by silver staining and streptavidin blotting, revealing strong and specific labeling of a band at 35-37 kDa that corresponds to the respective BSH monomers. We also verified that the labeled proteins are the expected BSHs from these bacteria using mass spectrometry-based proteomics (Data S2-S3). Therefore, BSH-TRAP using each of the four probes can pull down BSHs from *B. fragilis* and *B. longum*.

**Figure 5.**
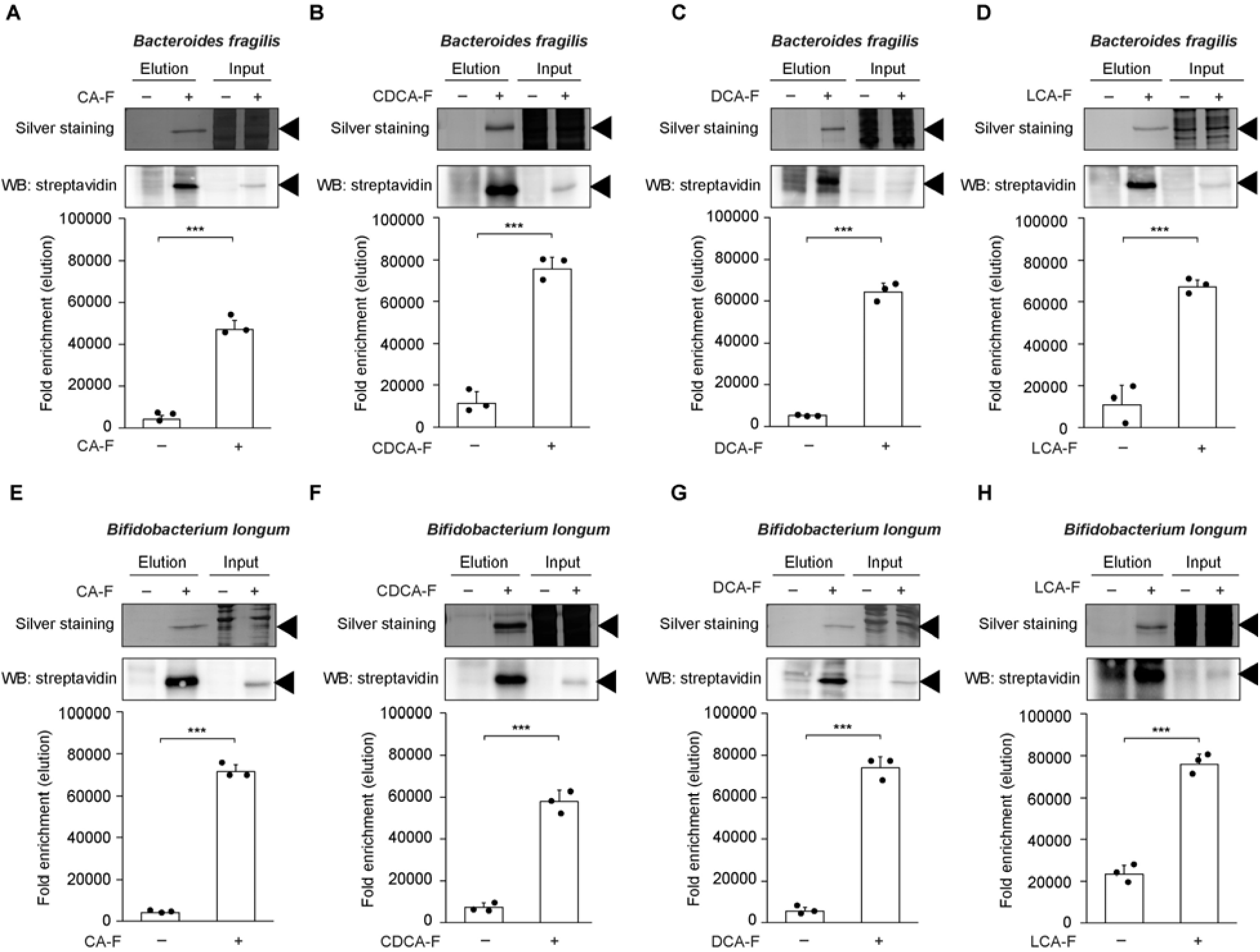
ABPs pull down BSH from human gut anaerobes. ABPs (indicated, 50 µM) were incubated with lysates from (A-D) *Bacteroides fragilis*, or (E-H) *Bifidobacterium longum* at 37°C for 1 h. CuAAC tagging was carried out with biotin-alkyne, followed by pulldown with streptavidin-agarose enrichment. Samples were subjected to SDS-PAGE and analyzed either by streptavidin-HRP blot or by silver staining. Input is 10% of the elution. Arrowhead indicates BSH monomers at 35-37 kDa. The bands were quantified by densitometry using ImageJ (bottom panels). Error bars represent standard deviation from the mean. One-way ANOVA followed by post hoc Tukey’s test: *** p<0.001, n = 3.

### ABPs can label and enrich active BSHs from the mouse gut microbiome

We then applied BSH-TRAP to the murine gut microbiome to determine if the approach could profile active BSHs within a more complex biological sample corresponding to a metaproteome from thousands of different microbes. Lysates from wild-type mouse fecal bacteria were treated with CA-F, CDCA-F, DCA-F, or LCA-F for 1 h, followed by CuAAC tagging with Fluor 647-alkyne and the in-gel fluorescence assay. We found that the panel of probes labeled active BSHs in mouse gut bacteria with expected molecular weights of each monomer ranging from 35-42 kDa, with no labeling observed when the samples were pre-treated with iodoacetamide (Figure 6A-D, Figure S5A-D). We also determined that the BSH ABPs could enable enrichment of active BSHs by treatment of the lysates with the BSH ABPs for 1 h, followed by CuAAC tagging with biotin-alkyne and enrichment with streptavidin agarose. The samples were analyzed by gel electrophoresis, followed by silver staining and streptavidin blotting, revealing strong labeling of proteins of a similar size range (Figure 6E-H, Figure S5E-H). These results reveal that BSH-TRAP can effectively pull down active BSHs from the mouse gut microbiome.

**Figure 6.**
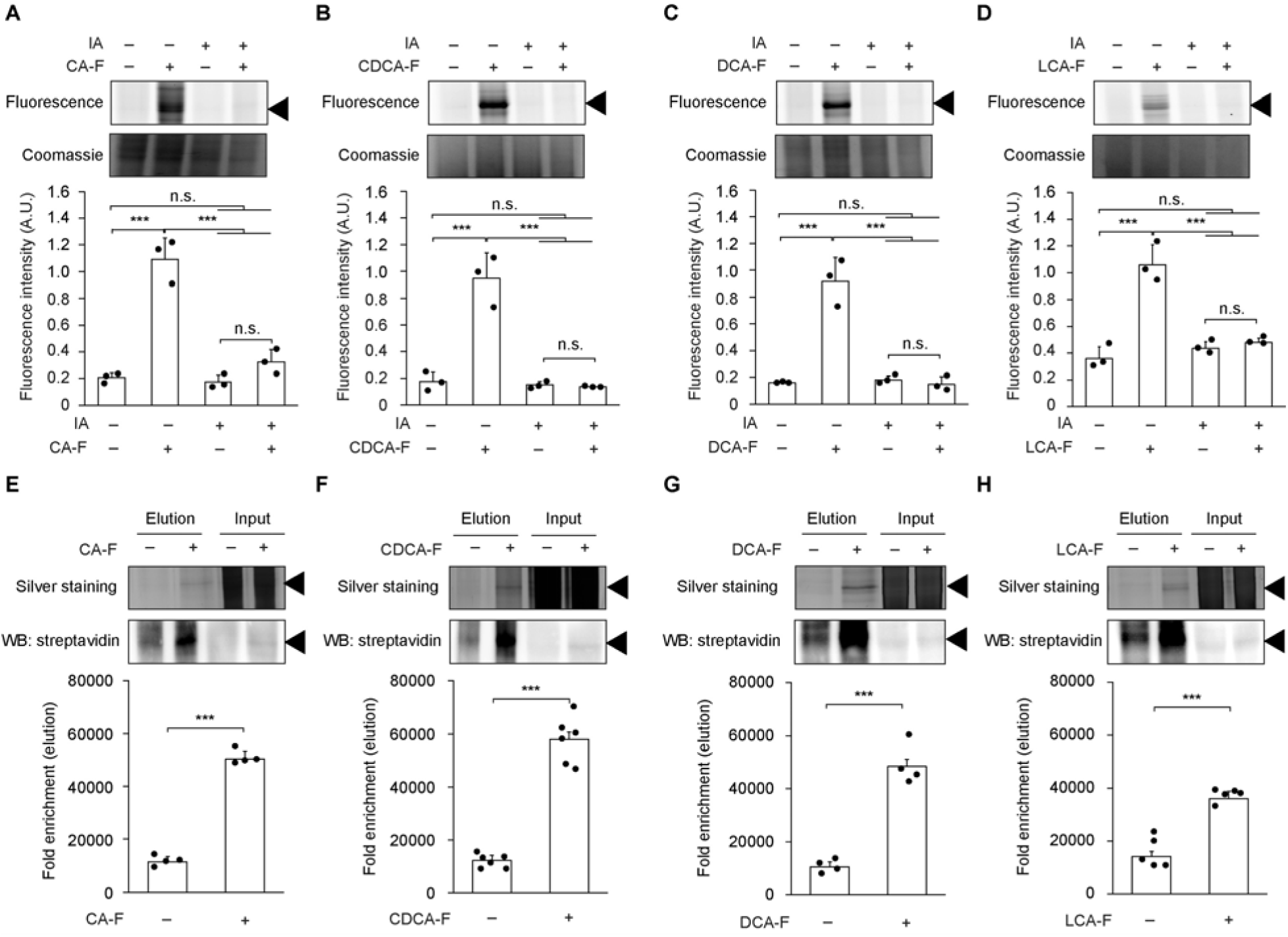
ABPs identify BSH activity in the mouse gut microbiome. (A-D) Bacterial lysates isolated from fecal pellets of healthy wild-type mice were incubated with ABPs (indicated, 50 µM) at 37 °C for 1 h. After CuAAC tagging with Fluor 647-alkyne, samples were analyzed by SDS-PAGE, followed by visualization using fluorescence. As a negative control for cysteine labeling, iodoacetamide (IA, 50 mM) was added prior to probe. Coomassie staining served as the loading control. (E-H) Alternatively, bacterial lysates were incubated with ABPs (indicated, 100 µM) at 37 °C for 1 h. After CuAAC tagging with biotin-alkyne, labeled proteins were enriched by streptavidin-agarose pulldown and analyzed either by streptavidin-HRP blot or by silver staining. Input is 10% of the elution. Arrowhead indicates 37 kDa ladder marker. The bands were quantified by densitometry using ImageJ (bottom panels). Error bars represent standard deviation from the mean. One-way ANOVA followed by post hoc Tukey’s test: *** p<0.001, n.s. = not significant, (A-D) n = 3, (E) n = 4, (F) n = 6, (G) n = 4, (H) n = 5.

### BSH-TRAP can profile BSH substrate specificity of the mouse gut microbiome

Finally, we applied BSH-TRAP to explore the substrate specificity of gut microbiota-associated BSHs within mice. To construct this metaproteome, we assembled the metagenome from the murine gut microbiome, including all *bsh* genes within our mouse colony and the Mouse Gastrointestinal Bacterial Catalogue (Beresford-Jones et al., 2022) (Data S4), and used Prodigal, a gene prediction program for microbial genomes (Hyatt et al., 2010), to create a protein database. Fecal bacterial lysates from wild-type mice were treated with CA-F, CDCA-F, DCA-F, and LCA-F for 1 h, followed by CuAAC tagging using biotin-alkyne. The labeled proteins were streptavidin-affinity enriched, and the samples were subjected to mass spectrometry-based proteomics using our metaproteomic database. As expected, all four probes enriched for BSHs over the vehicle control, and we verified their identities using shotgun proteomics (Figure S6A, Table 1, Data S5). These results demonstrate that BSH-TRAP enables the identification of active BSHs from the mouse gut microbiota metaproteome.

**Table 1.** Bacterial abbreviations.

| **Abbreviation** | **Taxonomic lineage** |
| --- | --- |
| *BE* | *Bacillota (phylum), Clostridia (class), Eubacteriales (order)* |
| *BM_1* | *Bacteroidota (phylum), Bacteroidia (class), Bacteroidales (order), Muribaculaceae (family)_1* |
| *BM_2* | *Bacteroidota (phylum), Bacteroidia (class), Bacteroidales (order), Muribaculaceae (family)_2* |
| *CC_1* | *Bacillota (phylum), Clostridia (class), Eubacteriales (order), Clostridiaceae (family), Clostridium (genus)_1* |
| *CC_2* | *Bacillota (phylum), Clostridia (class), Eubacteriales (order), Clostridiaceae (family), Clostridium (genus)_2* |
| *EA_1* | *Actinomycetota (phylum), Coriobacteriia (class), Eggerthellales (order), Eggerthellaceae (family), Adlercreutzia (genus)* |
| *EA_2* | *Actinomycetota (phylum), Coriobacteriia (class), Eggerthellales (order), Eggerthellaceae (family), Adlercreutzia (genus)* |
| *EF* | *Bacillota (phylum), Erysipelotrichia (class), Erysipelotrichales (order), Erysipelotrichaceae (family), Faecalibaculum (genus)* |
| *LJ_1* | *Bacillota (phylum), Bacilli (class), Lactobacillales (order), Lactobacillaceae (family), Lactobacillus (genus), Lactobacillus johnsonii (species)_1* |
| *LJ_2* | *Bacillota (phylum), Bacilli (class), Lactobacillales (order), Lactobacillaceae (family), Lactobacillus (genus), Lactobacillus johnsonii (species)_2* |
| *LJ_3* | *Bacillota (phylum), Bacilli (class), Lactobacillales (order), Lactobacillaceae (family), Lactobacillus (genus), Lactobacillus johnsonii (species)_3* |
| *LL* | *Bacillota (phylum), Bacilli (class), Lactobacillales (order), Lactobacillaceae (family), Limosilactobacillus (genus)* |
| *PF* | *Bacillota (phylum), Tissierellia (class), Tissierellales (order), Peptoniphilaceae (family), Finegoldia (genus)* |
| *PR* | *Bacillota* (phylum), *Clostridia* (class), *Eubacteriales* (order), *Peptostreptococcaceae* (family), *Romboutsia* (genus) |
| *TT_1* | *Bacillota* (phylum), *Erysipelotrichia* (class), *Erysipelotrichales* (order), *Turicibacteraceae* (family), *Turicibacter* (genus) |
| *TT_2* | *Bacillota* (phylum), *Erysipelotrichia* (class), *Erysipelotrichales* (order), *Turicibacteraceae* (family), *Turicibacter* (genus) |

Interestingly, these studies revealed that the BSHs that were enriched by our probes did not correspond to the most abundant bacterial *bsh* genes within the murine gut microbiome, as assessed by our metagenomic assemblies (Figure S6B-C, Data S6). Importantly, this finding illustrates that our chemoproteomic approach is not biased for highly abundant microbes and instead reveals enzymatic activity, which cannot be obtained by metagenomic sequencing alone. In these studies, we observed enrichment of a wide diversity of microbial BSHs, including those from the major bacterial phyla found in the gut, from both Gram-positive and Gram-negative bacteria, and from both facultative and obligate anaerobes (Figure S6A, Table 1). These findings demonstrate that BSH-TRAP can target a broad variety of gut microbiota-associated BSHs.

We then examined the substrate preference of the BSH from each bacterium that we were able to identify with high confidence from our metagenomic assemblies by comparing relative probe labeling, assuming that probe labeling correlates with BSH substrate preference for conjugated bile acids bearing that steroid core (Figure 7, Table 1; see also Figure 4). Interestingly, when we analyzed all of the pairwise comparisons of BSH labeling between two different probes, we found that each bacterial BSH exhibited distinct substrate preferences based on the bile acid steroid core. We further compared substrate preference across individual bacterial BSHs and found that CDCA was generally the most preferred bile acid substrate, whereas LCA was least preferred among the bacterial BSHs that we were able to confidently identify from our metagenomic assemblies (Figure 7, Table 1). However, careful analysis of this dataset revealed many subtleties of BSH substrate preference across the microbiome, as described in more detail in the Discussion section (vide infra). Thus, we have demonstrated that global BSHs activities within the mouse gut microbiome can be systemically profiled at the resolution of individual bacterial BSHs. These results illustrate the power of BSH-TRAP as a systems biochemistry approach that can simultaneously reveal substrate specificity of numerous proteins of the BSH enzyme family expressed by diverse members of the gut microbiota in a single systems-level study.

**Figure 7.**
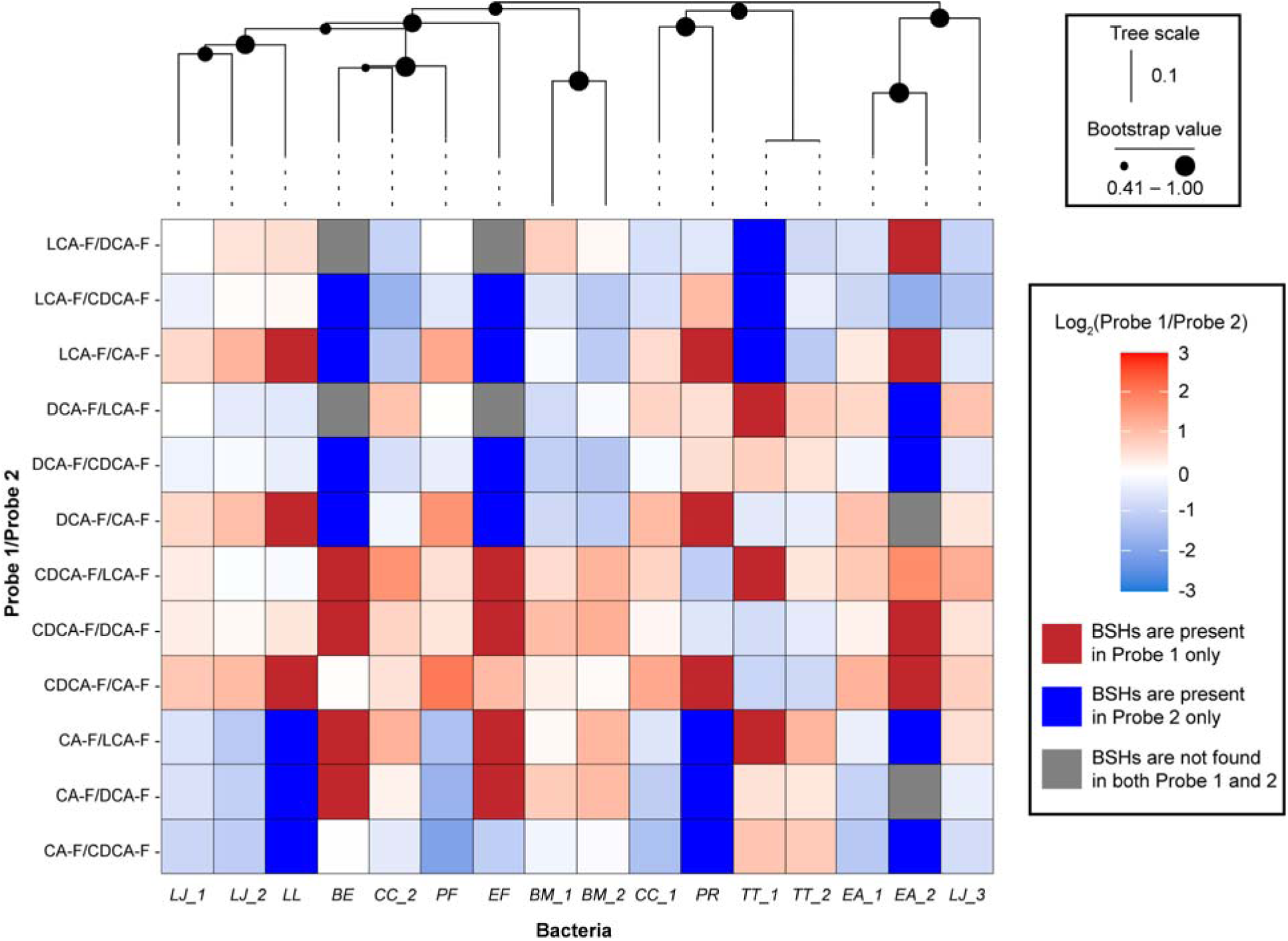
ABPs elucidate substrate specificity of gut microbiota-associated BSHs in the mouse gut microbiome. Heatmap of ABP-labeled BSHs identified by metagenomic assemblies. Bacterial lysates isolated from fecal pellets of healthy wild-type mice were incubated with ABPs (indicated, 100 µM) at 37 °C for 1 h. After CuAAC tagging with biotin-alkyne, labeled proteins were enriched by streptavidin-agarose pulldown and analyzed by mass spectrometry-based proteomics. Red indicates increased BSH labeling comparing probe 1 to probe 2, shown in log2 fold change according to heatmap (with dark red indicating labeling detected only with probe 1, i.e., absent in probe 2 samples), blue indicates decreased BSH labeling comparing probe 1 to probe 2 (with dark blue indicating labeling detected only with probe 2, i.e., absent in probe 1 samples), and gray indicates labeling not detected with both probes 1 and 2 (e.g., absent in probe 1 and 2 samples). Bacterial abbreviations are provided in Table 1. *N.B.*, When an abbreviation ends in “_N” where N=1, 2, etc., it indicates that more than one distinct bacteria was detected but because of the incomplete nature of shotgun metagenomic assembly that these different bacteria could not be fully distinguished at the taxonomic level. Pairwise distances from aligned sequences were calculated using Fitch matrix and used in neighbor-joining tree estimation (Fitch, 1966; Saitou and Nei, 1987; Studier and Keppler, 1998).

## Discussion

The metaproteome of the gut microbiome is a vastly complex biological system whose individual biochemical activities are challenging to probe. Given the astounding metabolic potential of the gut microbiota, it is critical to characterize the enzymes that carry out important biosynthetic transformations. We have developed a unique systems biochemistry approach, BSH-TRAP, employing ABPP to identify active BSHs within the gut microbiota. This chemoproteomic platform exploits tailored ABPs that target BSH, covalently label its active site cysteine residue, and contain a click chemistry handle that enables enrichment and identification of the active BSHs. Here, we employ BSH-TRAP using a panel of ABPs with various bile acid cores to determine the substrate specificity of BSHs within gut anaerobic bacteria and the mouse gut microbiome.

Building on our initial ABP for BSH-TRAP, an azido CA-based AOMK probe (Parasar et al., 2019), here we have elaborated this prototype into a comprehensive strategy by replacing AOMK with the more reactive, and therefore more sensitive, FMK warhead (Kam et al., 2004). In addition, we expanded the technology from simply CA to a panel of BSH ABPs based on four predominant human BAs, including CA, CDCA, DCA, and LCA (de Aguiar Vallim et al., 2013). First, we demonstrated that these ABPs label *C. perfringens* CGH recombinantly expressed by *E. coli* in a time- and concentration-dependent manner. We also determined that the probes were cysteine-selective because the labeling was eliminated in the C2S mutant. We next demonstrated that the panel of probes can detect endogenously expressed BSH in human gut anaerobes, *B. fragilis* and *B. longum*, as evidenced by strong labeling of a protein at the expected molecular weight for each bacterial BSH monomer that was eliminated in the presence of iodoacetamide. We also determined the BSH substrate specificity of these human gut bacteria and found that CA and LCA were the least favored substrates for *B. fragilis* and *B. longum*, respectively. To demonstrate that these second generation ABPs can pull down BSH within gut microbes, we enriched for active BSHs from *B. fragilis* and *B. longum* and verified their identities by mass spectrometry-based proteomics. Then, we employed BSH-TRAP to visualize BSH activity within bacterial lysates from the mouse gut microbiome and found that this complex biological sample contained multiple BSH monomers of different molecular weights (35-42 kDa), whose labeling can be competed away with iodoacetamide.

Finally, we applied BSH-TRAP using the panel of probes to profile the substrate specificity of active BSHs within mouse gut microbiota. Using metagenomics and chemoproteomics, we were able to confidently identify at least sixteen active BSHs from distinct bacteria within the murine gut microbiome with our panel of probes. Furthermore, our analysis of shotgun metagenomic assemblies of the mouse gut microbiota verified that the bacterial BSHs enriched by our panel of ABPs do not necessarily derive from the most abundant bacterial *bsh* genes based on quantification from our metagenomic assemblies. These results highlight the utility of our chemoproteomic approach in identifying enzyme activity, which cannot be predicted by metagenomics and proteomics alone.

In these studies, we found that although individual bacterial BSHs exhibited distinct substrate specificities based on the bile acid steroid core, bacteria that are more closely related by phylogenetic analysis of the BSH protein sequences had more similar substrate preferences (Figure 7, Table 1). For example, *Turicibacter* spp. preferred cholic acid-containing BAs, whereas *Muribaculaceae*, *Clostridium* spp., *Adlercreutzia* spp., and *Lactobacillus johnsonii* preferred chenodeoxycholic acid-containing substrates. The labeling of three BSHs from *L. johnsonii* enabled us to examine this phenomenon in more detail at the sub-species level. Interestingly, the two BSHs from *L. johnsonii* (*LJ_1* and *LJ_2*) that are 60.1% identical exhibited more similar substrate preferences. However, a third BSH from *L. johnsonii* (*LJ_3*) that shared less amino acid conservation at the protein sequence level (32.8% compared to *LJ_1* and 37.4% compared to *LJ_2*) exhibited a more distinct probe labeling profile, indicating different substrate preferences. These results are consistent with phylogenetic studies that suggest BSH protein sequence dictates substrate specificity for certain bacteria (Song et al., 2019).

In addition, our studies revealed that BSHs from certain bacteria had stronger substrate preferences (e.g., *BE*, *EA_2*, *EF*, *LL*, *PR*, and *TT_1*) due to their differential binding of the probes compared to BSHs from the remaining bacteria that were identified, suggesting that certain microbes may contribute more strongly to overall BSH activity. For instance, *LL* and *PR* strongly preferred CDCA, DCA, and LCA over CA, whereas *BE* and *EF* strongly preferred the two primary bile acids over the two secondary bile acids as substrates. Together, these results demonstrate that BSH-TRAP can enable profiling of global BSH activities within the gut microbiome at the systems level, while also identifying substrate preferences of individual bacterial BSHs. Moreover, these studies revealed that BSHs from individual gut bacteria exhibit differing substrate preferences based on the bile acid steroid core, suggesting that distinct BSHs contribute to the composition of the total bile acid pool within the host. Furthermore, the activities of certain bacterial BSHs on preferred conjugated bile acids may lead to activation of specific signaling pathways via bile acid receptors, including FXR and TGR5, that have strong preference for particular bile acid metabolites (Cai et al., 2022; Collins et al., 2023; Perino and Schoonjans, 2022).

In conclusion, systems biochemistry approaches such as BSH-TRAP can characterize enzymatic activities within complex biological systems, such as the metaproteome of the gut microbiome. We envision that similar chemical strategies using ABPP and tailored ABPs could be applied to characterize enzymes beyond BSHs to profile alternative metabolic pathways. In addition, application of ABPs targeting different enzymatic activities could reveal how global gut microbiota metabolism contributes to changes in levels of microbial metabolites within the host. Such approaches could also be exploited to elucidate substrate specificity within these biochemical pathways to understand how overall gut microbiota metabolism contributes to different physiological states. Finally, we expect that these chemoproteomic technologies could also reveal how differential enzymatic activities impact various pathological conditions and inflammatory diseases.

## Limitations of the study

One limitation of this study is that the design of our probes may preclude the ability to assess bile acid conjugate preference of BSHs (i.e., glyco-, tauro-, etc.), because our ABPs feature a covalent warhead whose structure differs substantially from the amino acid side chain that is at this position in the natural substrate. As a result, we believe that the probes for BSH-TRAP in this study can provide information on substrate specificity in circumstances where it is governed by the bile acid steroid core but not the amino acid side chain (Batta et al., 1984; Huijghebaert and Hofmann, 1986; Yao et al., 2018). As well, an inherent limitation of this approach is that metagenomics is necessary to construct the protein database used to identify murine gut bacteria with active BSHs, and metagenomic assembly remains a challenge in the field (New and Brito, 2020). Additionally, false negatives can arise from sample proteolysis because the proteomic analysis was limited to proteins of a specific size range that encompasses all known BSHs to reduce sample complexity. In addition, the ABPs used in this study were based on bile acid steroid cores that are the most abundant in humans, and their relative abundance differs from that in mice (de Aguiar Vallim et al., 2013). Therefore, we were only able to profile BSH substrate preference for the bile acid steroid cores included in our panel of probes. Consequently, the taxonomic identification of the active bacterial BSHs from the mouse gut microbiome represent the lower limit of BSHs that we could confidently report in this study due to the existence of false negatives. Finally, further biochemical characterization of individual BSHs identified in the proteomics study is necessary to distinguish between members of the linear amide C-N hydrolase, choloylglycine hydrolase family (Pfam PF02275), which include not only BSHs but also other hydrolases such as penicillin acylases.

## Significance

Bile acid metabolism is a complex process that is carried out by both the host and gut microbiota; however, the specific enzymes that perform these biotransformations remain difficult to characterize because of the sheer number of microbial enzymes within the intestines. Bile salt hydrolase (BSH) is considered the gatekeeper enzyme of secondary bile acid metabolism in the gut; yet, the substrate specificity of discrete members from this family within this complex ecosystem remains unknown. Here, we apply activity-based protein profiling (ABPP) using tailored covalent probes to profile BSH substrate specificity within the gut microbiota and find that individual BSHs from different bacteria prefer different bile acid substrates based on the steroid core. These results highlight the role of specific bacterial enzymes in determining bile acid composition within the gut.

## Acknowledgements

This work was supported by an NIH R35 Maximizing Investigators’ Research Award for Early Stage Investigators (R35GM133501). Research in the Chang Lab is supported by a Beckman Young Investigator Award (to P.V.C.) from the Arnold and Mabel Beckman Foundation and a Sloan Research Fellowship (to P.V.C.) from the Alfred P. Sloan Foundation. I.L.B. was supported by the Packard Foundation and the Pew Charitable Trusts. This work made use of the Cornell University NMR Facility, which is supported, in part, by the NSF through MRI award CHE-1531632. We thank the Proteomics and Metabolomics Facility of Cornell University for providing the mass spectrometry data and NIH SIG grant 1S10 OD017992-01 support for the Orbitrap Fusion mass spectrometer. We thank the Weill Institute for Cell and Molecular Biology for additional resources. We also thank N. Frederick and R. Vignogna for technical assistance.

## Author Contributions

Conceptualization, L.H. and P.V.C.; investigation, L.H., A.P., A.S., R.X., S.A.S., J.A.P., K.P.M., and P.D.; methodology, L.H. and P.V.C.; formal analysis, L.H., A.P., and A.S.; visualization, L.H.; writing – original draft, L.H., I.L.B., and P.V.C.; writing – review & editing, L.H., I.L.B., and P.V.C.; project administration, P.V.C.; supervision, I.L.B. and P.V.C.; funding acquisition, I.L.B. and P.V.C.

## Inclusion and Diversity

We support inclusive, diverse, and equitable conduct of research.

## Supplemental information titles and legends

**Figure S1. Activity-based probes (ABPs) label active *Clostridium perfringens* choloylglycine hydrolase (CGH).** Wild-type or point mutant (Cys2Ser, C2S) CGH (1.5 µg) from *C. perfringens* was treated with 10 µM of (A) CA-F, (C) CDCA-F, (E) DCA-F, and (G) LCA-F for various amounts of time (indicated) at 37 °C. Alternatively, wild-type *C. perfringens* CGH (1.5 µg) was treated with (B) CA-F, (D) CDCA-F, (F) DCA-F, and (H) LCA-F in a dose-dependent manner (indicated) for 1 h at 37 °C. The samples were labeled using CuAAC tagging with Fluor 488-alkyne. The samples were purified by SDS-PAGE and visualized by fluorescence (excitation wavelength = 488 nm). The gel was stained with Coomassie brilliant blue as a loading control (added bovine serum albumin appears at ∼66 kDa). Arrowhead indicates CGH monomer at 37 kDa.

**Figure S2. ABPs label bile salt hydrolases (BSHs) in human gut anaerobes.** (A-D) *Bacteroides fragilis* lysate was treated with 10 µM of (A) CA-F, (B) CDCA-F, (C) DCA-F, and (D) LCA-F for 1 h at 37 °C. (E-H) *Bifidobacterium longum* lysate was labeled with 10 µM of (E) CA-F, (F) CDCA-F, (G) DCA-F, and (H) LCA-F for 1 h at 37 °C. Lysates were treated with or without 50 mM of iodoacetamide (IA) prior to incubation with ABPs as a negative control. Following probe labeling, CuAAC tagging was carried out with Fluor 488-alkyne, and samples were subjected to SDS-PAGE. The gel was visualized using fluorescence, followed by staining with Coomassie brilliant blue. Arrowhead indicates BSH monomers at 35-37 kDa.

**Figure S3. ABPs reveal BSH substrate specificity in *B. fragilis* and *B. longum.*** Lysates from (A) *B. fragilis* and (B) *B. longum* were treated with CA-F, CDCA-F, DCA-F, or LCA-F (50 µM) for 10 min at 37 °C. Following probe labeling, CuAAC tagging was carried out with Fluor 488-alkyne, and samples were subjected to SDS-PAGE. The gel was visualized using fluorescence, followed by staining with Coomassie brilliant blue. Arrowhead indicates BSH monomers at 35-37 kDa.

**Figure S4. ABPs enrich BSHs in human gut anaerobes**. (A-D) *B. fragilis* lysates were treated with (A) CA-F, (B) CDCA-F, (C) DCA-F, and (D) LCA-F (50 µM) for 1 h at 37 °C. (E-H) *B. longum* lysates were treated with (E) CA-F, (F) CDCA-F, (G) DCA-F, and (H) LCA-F (50 µM) for 1 h at 37 °C. The labeled proteins were tagged with biotin-alkyne using CuAAC, enriched by streptavidin pulldown, and then purified by SDS-PAGE, followed by analysis by Western blot with streptavidin-HRP or silver staining. Input is 10% of the elution. Arrowhead indicates BSH monomers at 35-37 kDa.

**Figure S5. ABPs label endogenous BSHs in bacteria from the mouse gut microbiome.** (A-D) Lysates from mouse fecal bacteria were treated with (A) CA-F, (B) CDCA-F, (C) DCA-F, and (D) LCA-F (50 µM) for 1 h at 37 °C. Samples were treated with or without 50 mM of iodoacetamide (IA) prior to incubation with probes. The labeled proteins were tagged with Fluor 647-alkyne using CuAAC, and then purified by SDS-PAGE and visualized by fluorescence. The gel was stained with Coomassie brilliant blue as a loading control. (E-H) Lysates from mouse fecal bacteria were treated with (E) CA-F, (F) CDCA-F, (G) DCA-F, and (H) LCA-F (100 µM) for 1 h at 37 °C. The labeled proteins were tagged with biotin-alkyne using CuAAC, enriched by streptavidin pulldown, and then purified by SDS-PAGE, followed by analysis by Western blot with streptavidin-HRP or silver staining. Input is 10% of the elution. Arrowhead indicates 37 kDa ladder marker.

**Figure S6. ABPs identify bacterial BSHs in the mouse gut microbiome.** Lysates from mouse fecal bacteria were treated with CA-F, CDCA-F, DCA-F, and LCA-F (100 µM) for 1 h at 37 °C. The labeled proteins were tagged with biotin-alkyne using CuAAC, enriched by streptavidin pulldown, and analyzed by mass spectrometry-based proteomics. (A) Volcano plot of proteins that were detected in at least two replicate samples. The x-axis shows the log_2_(fold change), where fold change represents the ratio of protein abundance in the probe vs control samples, and the y-axis shows the −log_10_(P value), n= 3. Bacterial abbreviations are provided in Table 1. (B) Fold change (log_2_) of enrichment of BSHs from mouse gut microbiomes comparing probe treatment to control samples (y-axis) versus *bsh* gene abundance using reads per kilobase million (RPKM) in the mouse metagenomic assemblies (x-axis). (C) Relative abundance of phyla from the mouse metagenome based on bacterial taxonomic classification.

## TOC graphic

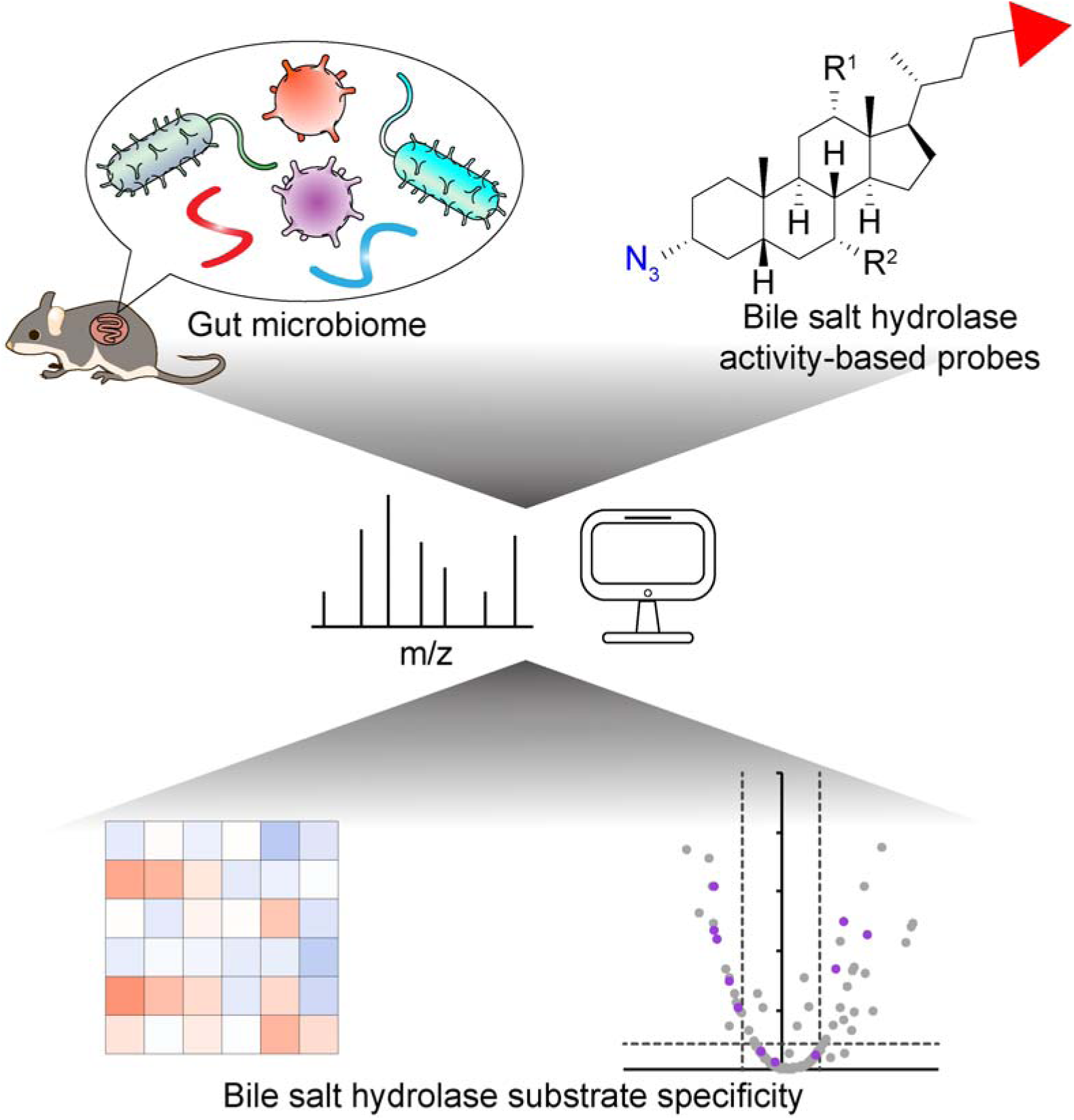

