## Supplementary Information for "Chemoproteomic profiling of substrate specificity in gut microbiota-associated bile salt hydrolases"

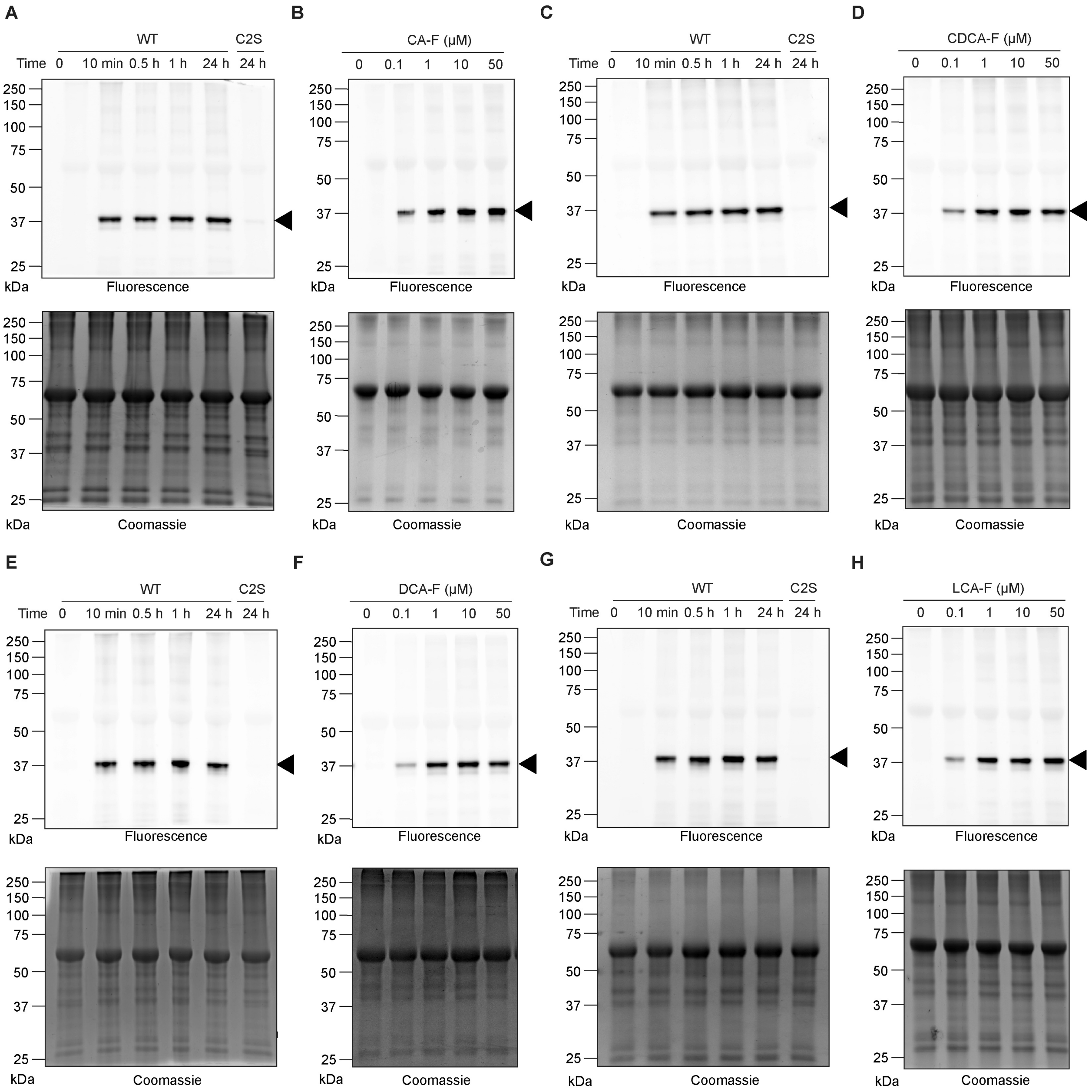
A
WT
C2S
Time 0 10 min 0.5 h 1 h 24 h 24 h
250
150
100
75
50
37
25
kDa
Fluorescence
Coomassie
B
CA-F (μM)
0 0.1 1 10 50
C
D
CDCA-F (μM)
E
F
DCA-F (μM)
G
H
LCA-F (μM)

**Figure S1**. **Activity-based probes (ABPs) label active *Clostridium perfringens* choloylglycine hydrolase (CGH).** Wild-type or point mutant (Cys2Ser, C2S) CGH (1.5 µg) from *C. perfringens* was treated with 10 µM of (A) CA-F, (C) CDCA-F, (E) DCA-F, and (G) LCA-F for various amounts of time (indicated) at 37 °C. Alternatively, wild-type *C. perfringens* CGH (1.5 µg) was treated with (B) CA-F, (D) CDCA-F, (F) DCA-F, and (H) LCA-F in a dose-dependent manner (indicated) for 1 h at 37 °C. The samples were labeled using CuAAC tagging with Fluor 488-alkyne. The samples were purified by SDS-PAGE and visualized by fluorescence (excitation wavelength = 488 nm). The gel was stained with Coomassie brilliant blue as a loading control (added bovine serum albumin appears at ~66 kDa). Arrowhead indicates CGH monomer at 37 kDa.

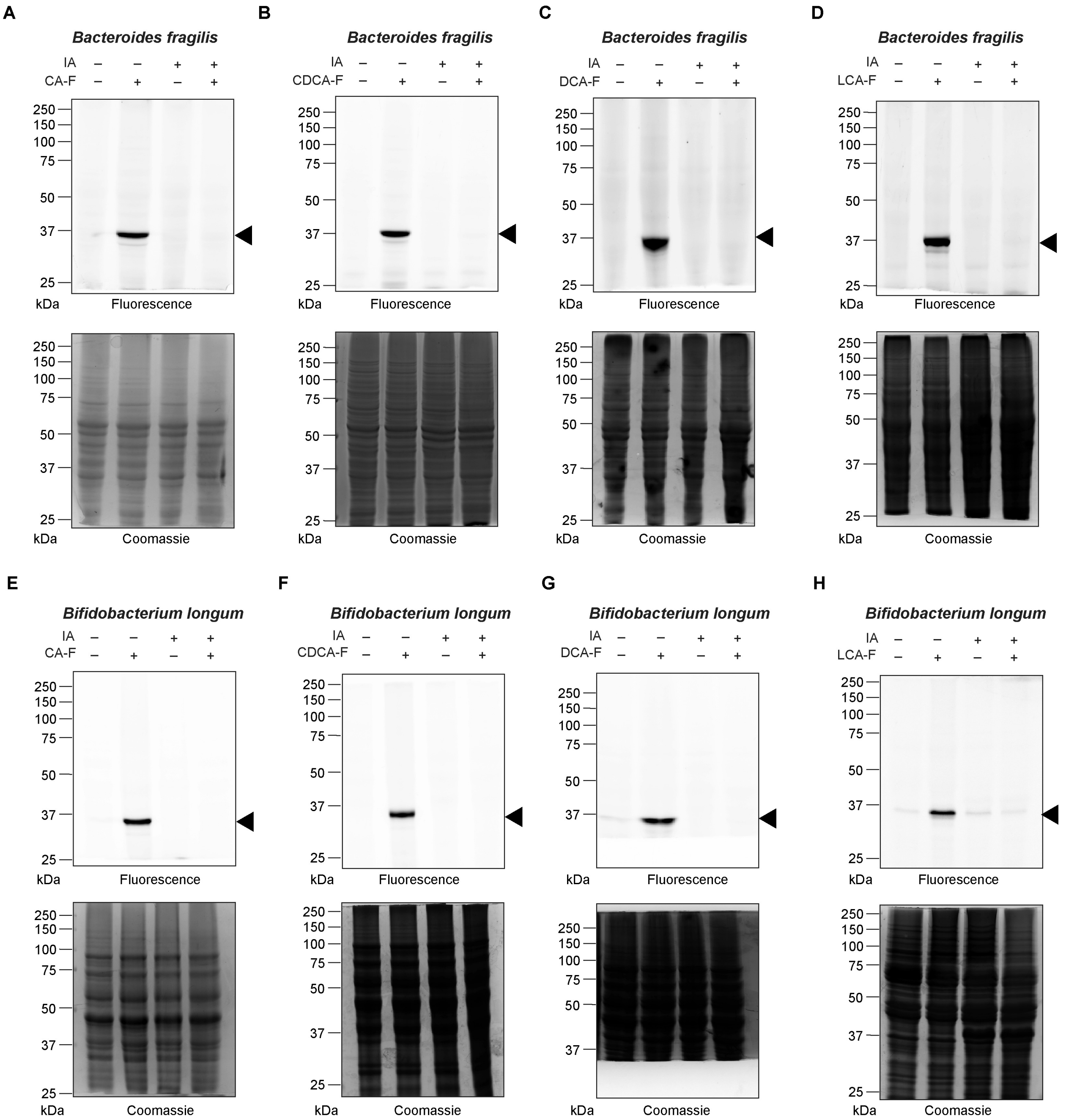
A
Bacteroides fragilis
IA – – + +
CA-F – + – +
250
150
100
75
50
37
25
kDa
Fluorescence
Coomassie
B
Bacteroides fragilis
IA – – + +
CDCA-F – + – +
Fluorescence
Coomassie
C
Bacteroides fragilis
IA – – + +
DCA-F – + – +
Fluorescence
Coomassie
D
Bacteroides fragilis
IA – – + +
LCA-F – + – +
Fluorescence
Coomassie
E
Bifidobacterium longum
IA – – + +
CA-F – + – +
Fluorescence
Coomassie
F
Bifidobacterium longum
IA – – + +
CDCA-F – + – +
Fluorescence
Coomassie
G
Bifidobacterium longum
IA – – + +
DCA-F – + – +
Fluorescence
Coomassie
H
Bifidobacterium longum
IA – – + +
LCA-F – + – +
Fluorescence
Coomassie

**Figure S2**. **ABPs label bile salt hydrolases (BSHs) in human gut anaerobes.** (A-D) *Bacteroides fragilis* lysate was treated with 10 μM of (A) CA-F, (B) CDCA-F, (C) DCA-F, and (D) LCA-F for 1 h at 37 °C. (E-H) *Bifidobacterium longum* lysate was labeled with 10 μM of (E) CA-F, (F) CDCA-F, (G) DCA-F, and (H) LCA-F for 1 h at 37 °C. Lysates were treated with or without 50 mM of iodoacetamide (IA) prior to incubation with ABPs as a negative control. Following probe labeling, CuAAC tagging was carried out with Fluor 488-alkyne, and samples were subjected to SDS-PAGE. The gel was visualized using fluorescence, followed by staining with Coomassie brilliant blue. Arrowhead indicates BSH monomers at 35-37 kDa.

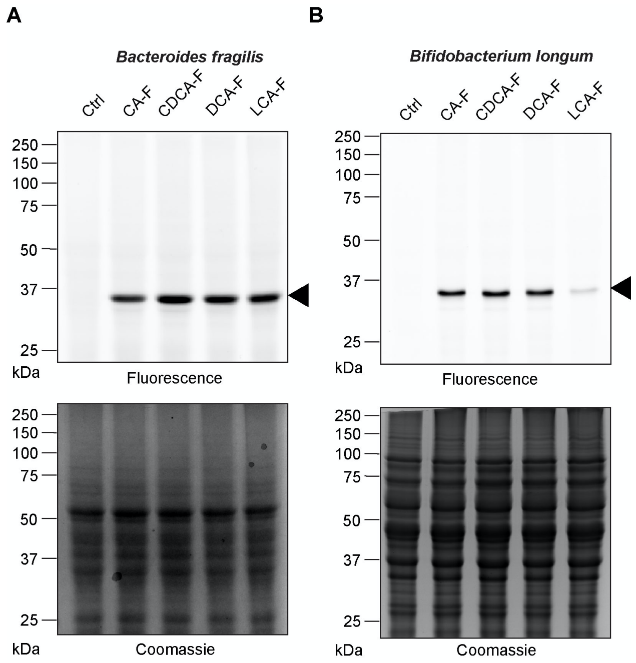


**Figure S3**. **ABPs reveal BSH substrate specificity in *B. fragilis* and *B. longum.*** Lysates from (A) *B. fragilis* and (B) *B. longum* were treated with CA-F, CDCA-F, DCA-F, or LCA-F (50 µM) for 10 min at 37 °C. Following probe labeling, CuAAC tagging was carried out with Fluor 488-alkyne, and samples were subjected to SDS-PAGE. The gel was visualized using fluorescence, followed by staining with Coomassie brilliant blue. Arrowhead indicates BSH monomers at 35-37 kDa.

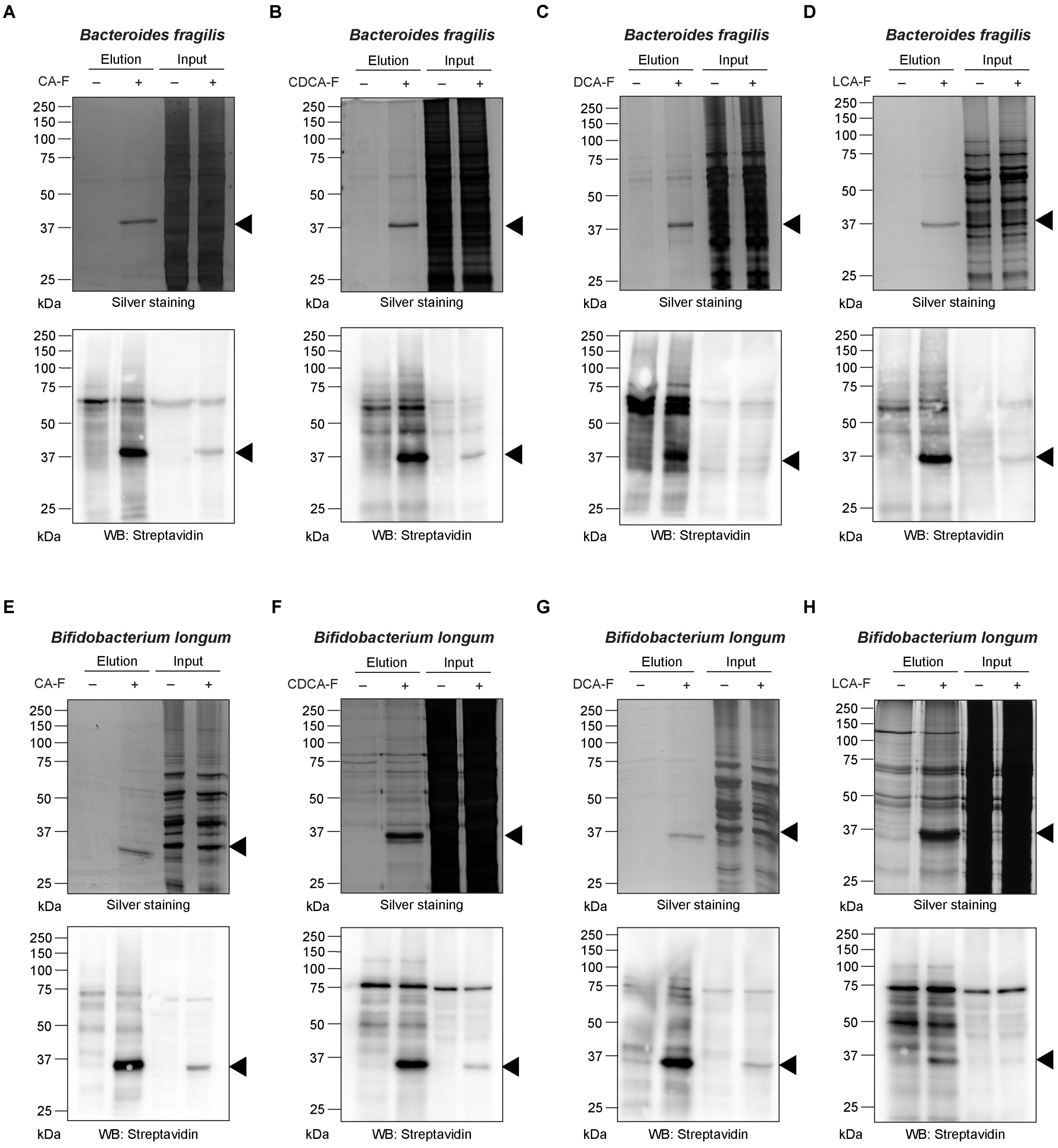


**Figure S4**. **ABPs enrich BSHs in human gut anaerobes**. (A-D) *B. fragilis* lysates were treated with (A) CA-F, (B) CDCA-F, (C) DCA-F, and (D) LCA-F (50 μM) for 1 h at 37 °C. (E-H) *B. longum* lysates were treated with (E) CA-F, (F) CDCA-F, (G) DCA-F, and (H) LCA-F (50 μM) for 1 h at 37 °C. The labeled proteins were tagged with biotin-alkyne using CuAAC, enriched by streptavidin pulldown, and then purified by SDS-PAGE, followed by analysis by Western blot with streptavidin-HRP or silver staining. Input is 10% of the elution. Arrowhead indicates BSH monomers at 35-37 kDa.

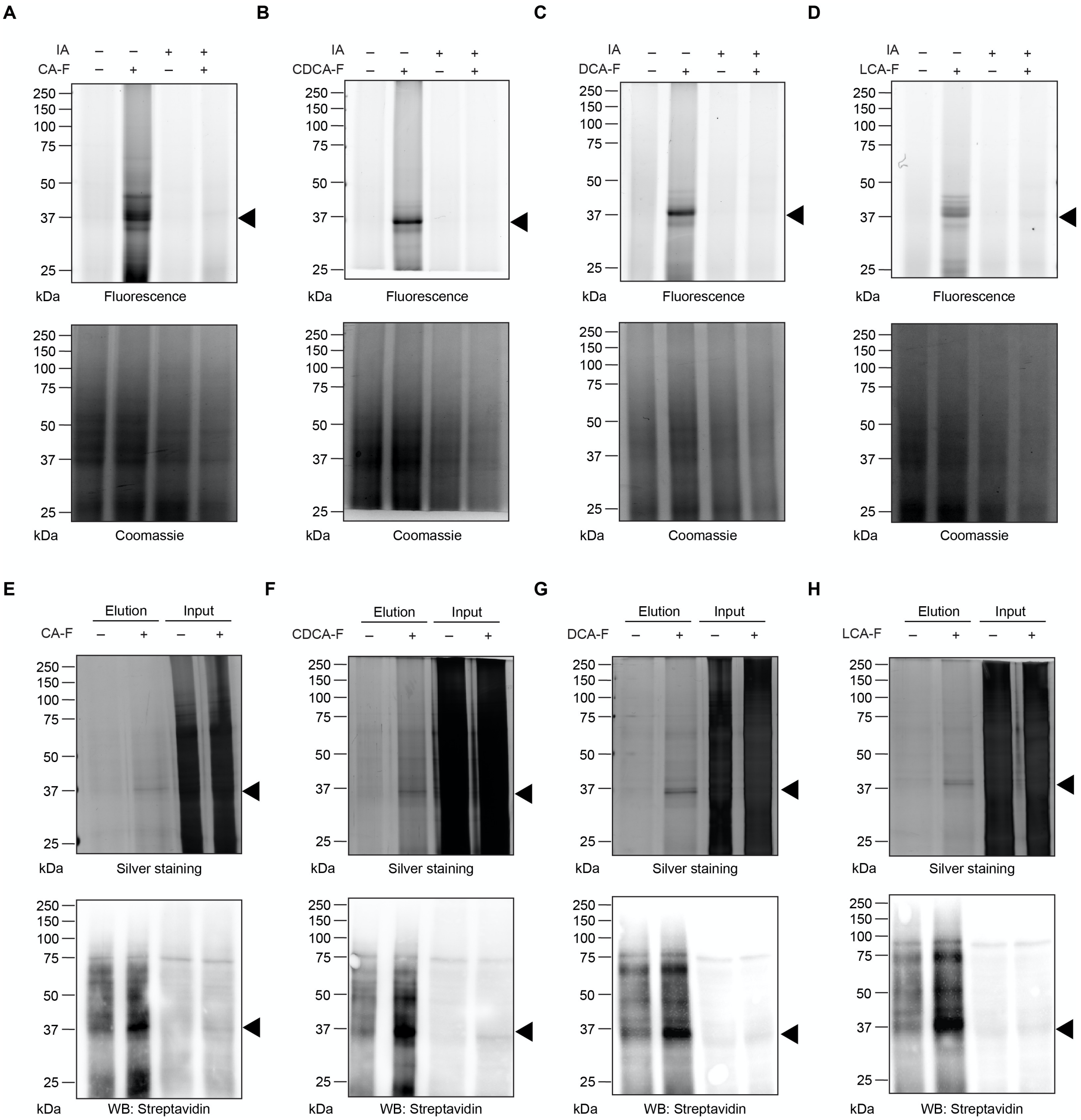


**Figure S5**. **ABPs label endogenous BSHs in bacteria from the mouse gut microbiome.** (A-D) Lysates from mouse fecal bacteria were treated with (A) CA-F, (B) CDCA-F, (C) DCA-F, and (D) LCA-F (50 μM) for 1 h at 37 °C. Samples were treated with or without 50 mM of iodoacetamide (IA) prior to incubation with probes. The labeled proteins were tagged with Fluor 647-alkyne using CuAAC, and then purified by SDS-PAGE and visualized by fluorescence. The gel was stained with Coomassie brilliant blue as a loading control. (E-H) Lysates from mouse fecal bacteria were treated with (E) CA-F, (F) CDCA-F, (G) DCA-F, and (H) LCA-F (100 μM) for 1 h at 37 °C. The labeled proteins were tagged with biotin-alkyne using CuAAC, enriched by streptavidin pulldown, and then purified by SDS-PAGE, followed by analysis by Western blot with streptavidin-HRP or silver staining. Input is 10% of the elution. Arrowhead indicates 37 kDa ladder marker.

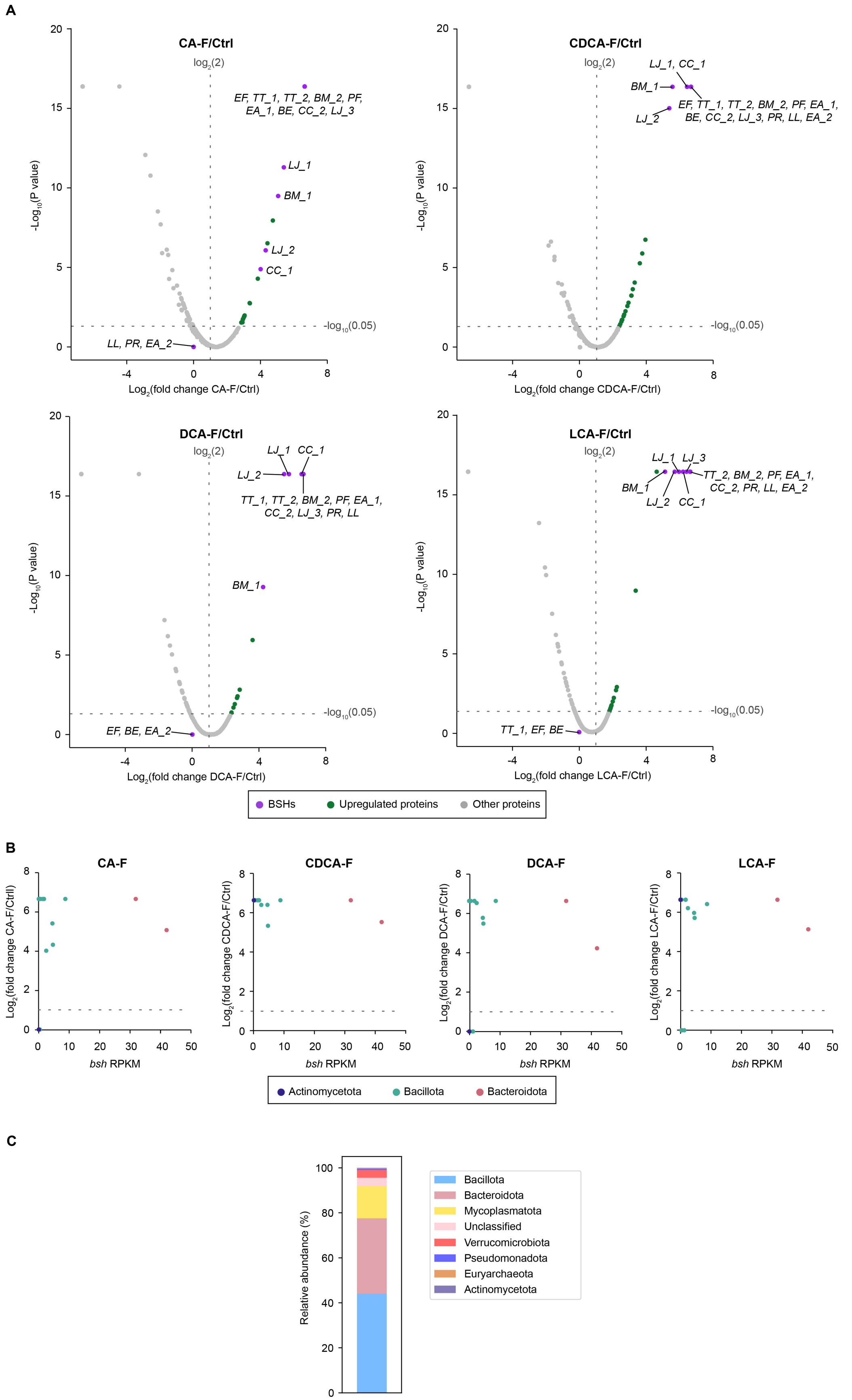
A
CA-F/Ctrl
log2(2)
-log10(0.05)
EF, TT_1, TT_2, BM_2, PF, EA_1, BE, CC_2, LJ_3
LJ_1
BM_1
LJ_2
CC_1
LL, PR, EA_2
-Log10(P value)
Log2(fold change CA-F/Ctrl)
CDCA-F/Ctrl
LJ_1, CC_1
BM_1
LJ_2
EF, TT_1, TT_2, BM_2, PF, EA_1, BE, CC_2, LJ_3, PR, LL, EA_2
Log2(fold change CDCA-F/Ctrl)
DCA-F/Ctrl
LJ_1
CC_1
LJ_2
TT_1, TT_2, BM_2, PF, EA_1, CC_2, LJ_3, PR, LL
BM_1
EF, BE, EA_2
Log2(fold change DCA-F/Ctrl)
LCA-F/Ctrl
LJ_1
LJ_3
TT_2, BM_2, PF, EA_1, CC_2, PR, LL, EA_2
BM_1
LJ_2
CC_1
TT_1, EF, BE
Log2(fold change LCA-F/Ctrl)
BSHs
Upregulated proteins
Other proteins
B
CA-F
CDCA-F
DCA-F
LCA-F
Log2(fold change CA-F/Ctrl)
Log2(fold change CDCA-F/Ctrl)
Log2(fold change DCA-F/Ctrl)
Log2(fold change LCA-F/Ctrl)
bsh RPKM
Actinomycetota
Bacillota
Bacteroidota
C
Relative abundance (%)
Bacillota
Bacteroidota
Mycoplasmatota
Unclassified
Verrucomicrobiota
Pseudomonadota
Euryarchaeota
Actinomycetota

**Figure S6**. **ABPs identify bacterial BSHs in the mouse gut microbiome.** Lysates from mouse fecal bacteria were treated with CA-F, CDCA-F, DCA-F, and LCA-F (100 μM) for 1 h at 37 °C. The labeled proteins were tagged with biotin-alkyne using CuAAC, enriched by streptavidin pulldown, and analyzed by mass spectrometry-based proteomics. (A) Volcano plot of proteins that were detected in at least two replicate samples. The x-axis shows the $\log_2$(fold change), where fold change represents the ratio of protein abundance in the probe (indicated) vs control (Ctrl) samples, and the y-axis shows the $-\log_{10}$(P value), n= 3. Bacterial abbreviations are provided in Table 1. (B) Fold change ($\log_2$) of enrichment of BSHs from mouse gut microbiomes comparing probe treatment to control samples (y-axis) versus *bsh* gene abundance using reads per kilobase million (RPKM) in the mouse metagenomic assemblies (x-axis). (C) Relative abundance of phyla from the mouse metagenome based on bacterial taxonomic classification.
